# Exponential growth of the USA overdose epidemic

**DOI:** 10.1101/134403

**Authors:** Jeanine M. Buchanich, Lauren C. Balmert, Donald S. Burke

**Affiliations:** Departments of Biostatistics, Graduate School of Public Health, University of Pittsburgh Pittsburgh, Pennsylvania, USA; Departments of Epidemiology, Graduate School of Public Health, University of Pittsburgh Pittsburgh, Pennsylvania, USA; Departments of Public Health Dynamics Laboratory, Graduate School of Public Health, University of Pittsburgh Pittsburgh, Pennsylvania, USA

## Introduction

The USA is experiencing an epidemic of drug overdose deaths ^1-3^. This epidemic has been attributed in part to increases in opioid prescribing that began in the late 1990s, which led to increasing substance use disorders and overdose deaths^4-7^. In an effort to understand and forecast trends in the epidemic, we examined data on reported accidental poisoning deaths from 1979 through 2015. We present our preliminary findings here.

## Methods

All reported accidental poisoning deaths were abstracted for the period 1979 to 2015 from the death record reports provided by the US National Center for Health Statistics and maintained in the MOIRA death record repository at the University of Pittsburgh. Cases were identified as those carrying codes for accidental poisoning in the 9^th^ or 10^th^ revisions of the International Classification of Diseases (ICD-9 codes E850 to E858 for 1979-1998, and ICD-10 codes X40 to X44 for 1999-2015). These codes included all drug poisonings but excluded alcohol and organic solvents and halogenated hydrocarbons and their vapors, other gases and vapors, pesticides, unspecified chemicals and noxious substances. For the purpose of this report we refer to the included accidental poisoning deaths collectively as drug overdose deaths. Population counts and age distributions were abstracted from the US Census. Overdose deaths per year were plotted on linear and logarithmic scales for the entire USA, and for all 50 states and the District of Columbia, and linear models were fit to the resulting data plots.

More than one half million overdose deaths were reported in the USA from 1979 through 2015 (Table 1). When deaths per year are plotted, the resultant curve appears to be exponential (Figure 1). A log transformed plot of the data shows that a simple exponential curve

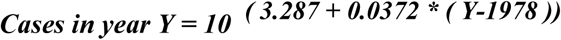

**Figure 1.**
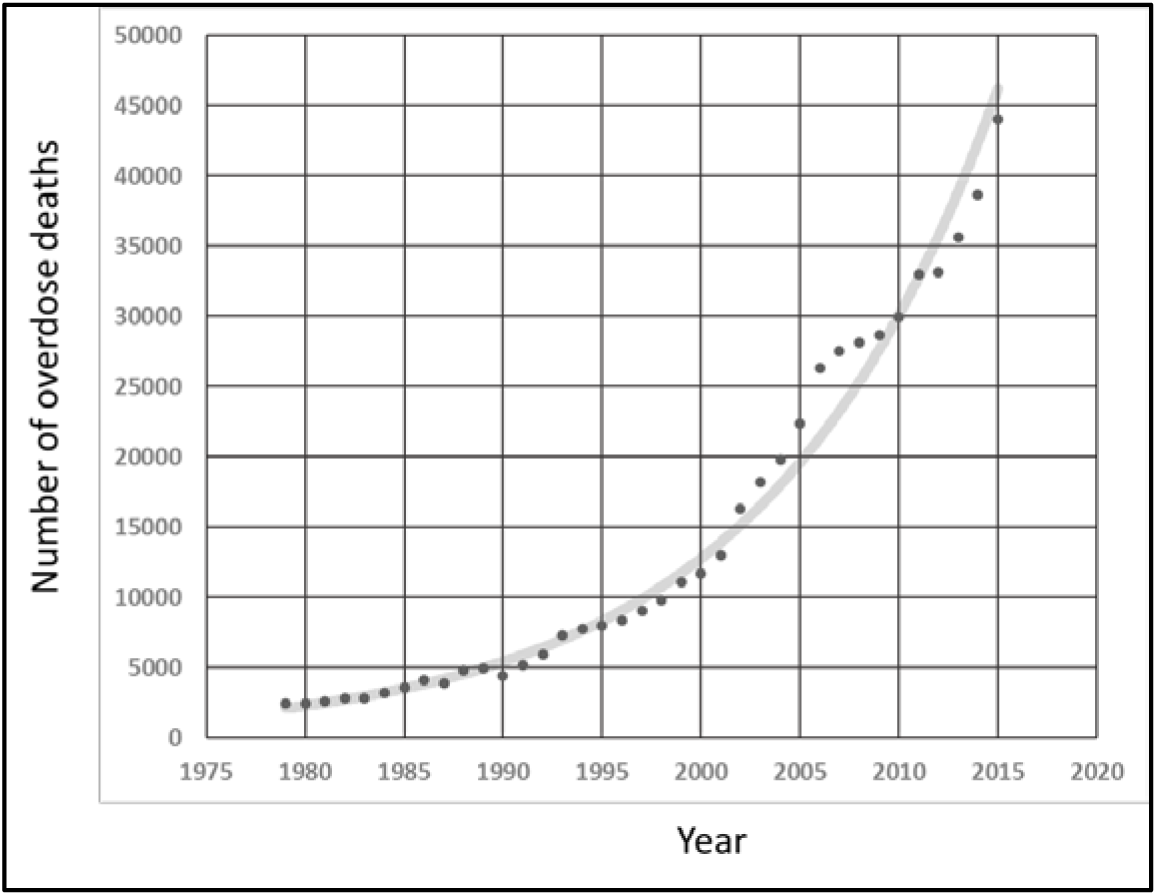
*Graph of the total number of drug overdose deaths per year in the USA from 1979 to 2015, including all ages, sexes, races, locations, and drugs. The plot appears to show exponential growth.*

**Table 1.**
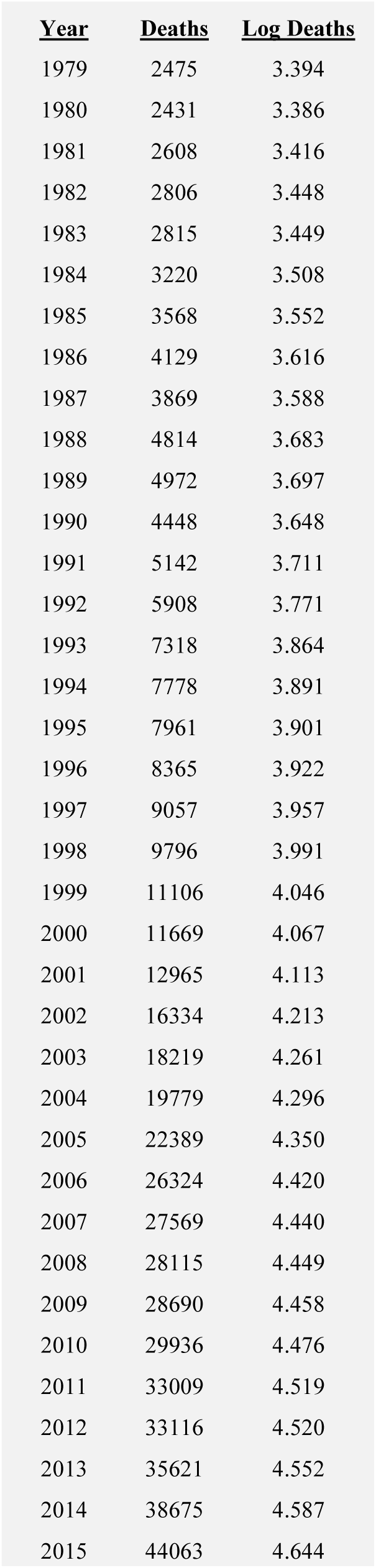
Results.

fits these plotted annual death counts with an R^2^ = 0.99 (Figure 2). Restated, total overdose death counts per year in the USA have increased on a nearly perfect exponential curve for at least the last 37 years. The average increase in deaths since 1979 has been 9% per year with an approximately 8 year doubling time. There have been some minor deviations from a perfect exponential trajectory, for example, a transient acceleration of the epidemic from 2002 to 2006, followed by a transient deceleration in 2007 to 2009, but overall the annual counts regularly return to a pattern of exponential growth. This pattern cannot be attributed to population growth, because annual US population increases since 1946 have never exceed 2.1% per year. Furthermore, the age-adjusted drug overdose death rates per 100,000 population also tightly follow exponential curves, for both females and males (R^2^ = 0.94 and R^2^ = 0.99, respectively, data not shown). Extrapolation of the 37 year log-linear plots of absolute death counts per year for five years into the future (2016 - 2020) forecasts that 300,089 additional drug overdose deaths will occur in the USA over that period if the trend is not deflected (Figure 2). Note that we include a “forecast” for the year 2016 because the National Center for Health Statistics Death Reports data for this most recent year are not yet available.

**Figure 2.**
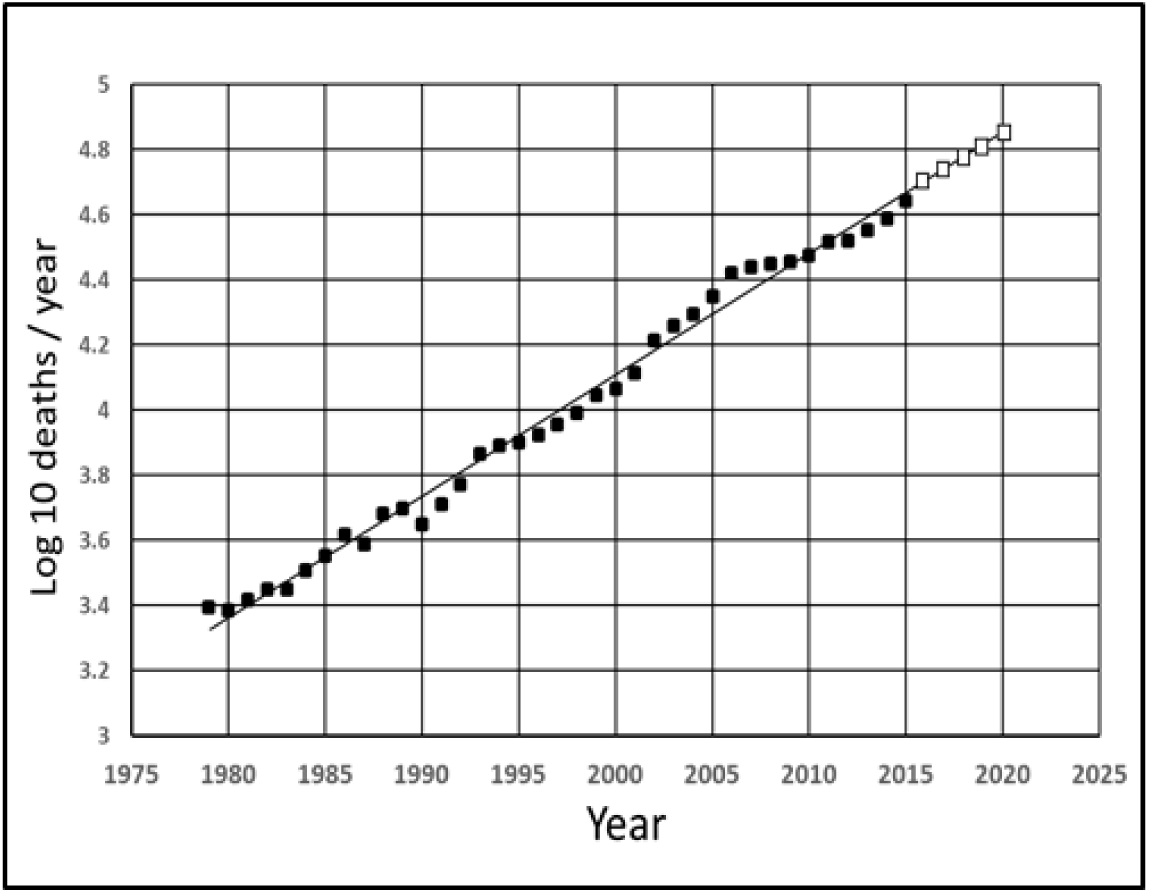
*Same data as in Figure 1 replotted as the log base 10 of the total number of drug overdose deaths per year. The plot is linear (R^2^ = 0.99). The fitted line can be extrapolated (empty squares) to forecast the expected number of deaths in the next five years (50,199 + 54,689 + 59579 + 64908 + 70713 = 300,089)*

Analyses of overdose death count data by state reveals that, like the national pattern, 45 states have experienced a clear exponential growth pattern (R^2^ > 0.8). For North Dakota, South Dakota, District of Columbia, Massachusetts, Maryland, and Rhode Island a simple exponential curve provides a looser fit to the observed data (R^2^ < 0.8). Extrapolations of the exponential growth pattern for each state can be employed to make state-specific forecasts (Table 2).

**Table 2.**
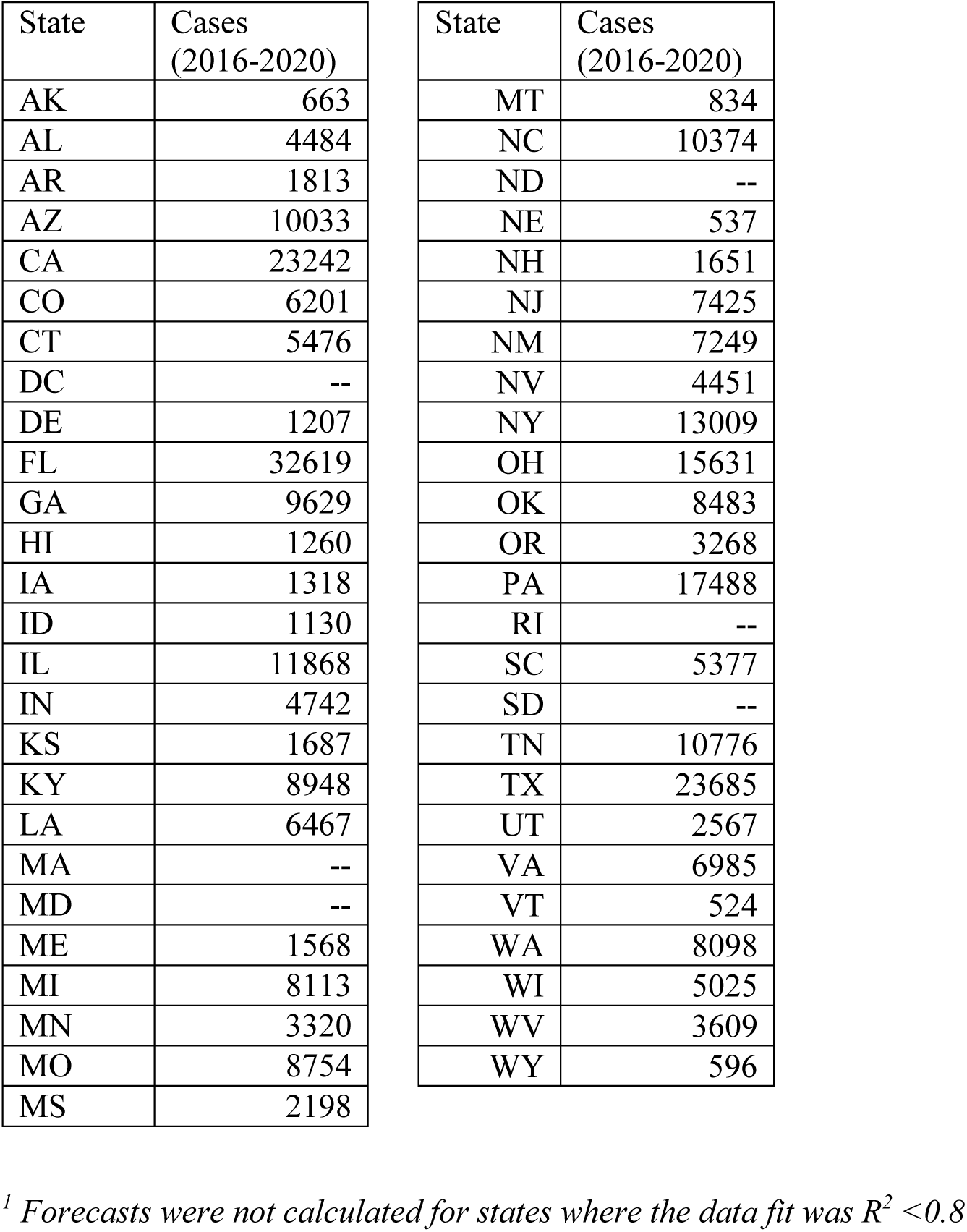
Forecasts of Accidental Poisoning Deaths by State Using Extrapolations from 1999-2015^1^

| State | Cases<br>(2016-2020) | State | Cases<br>(2016-2020) |
| --- | --- | --- | --- |
| AK | 663 | MT | 834 |
| AL | 4484 | NC | 10374 |
| AR | 1813 | ND | -- |
| AZ | 10033 | NE | 537 |
| CA | 23242 | NH | 1651 |
| CO | 6201 | NJ | 7425 |
| CT | 5476 | NM | 7249 |
| DC | -- | NV | 4451 |
| DE | 1207 | NY | 13009 |
| FL | 32619 | OH | 15631 |
| GA | 9629 | OK | 8483 |
| HI | 1260 | OR | 3268 |
| IA | 1318 | PA | 17488 |
| ID | 1130 | RI | -- |
| IL | 11868 | SC | 5377 |
| IN | 4742 | SD | -- |
| KS | 1687 | TN | 10776 |
| KY | 8948 | TX | 23685 |
| LA | 6467 | UT | 2567 |
| MA | -- | VA | 6985 |
| MD | -- | VT | 524 |
| ME | 1568 | WA | 8098 |
| MI | 8113 | WI | 5025 |
| MN | 3320 | WV | 3609 |
| MO | 8754 | WY | 596 |
| MS | 2198 |  |  |

<sup>1</sup> Forecasts were not calculated for states where the data fit was $R^2 < 0.8$

## Discussion

Analysis of death record reports shows that drug overdose deaths have been increasing exponentially in the USA since 1979. This exponential growth may have been occurring longer, but data prior to 1979 are difficult to interpret because of coding changes that occurred between the 8^th^ and 9^th^ revisions of the International Classification of Diseases. An exponential epidemic growth pattern is also observed at the state level for most states. Preliminary analyses (data not shown) suggest that there has been an exponential growth pattern of the epidemic in most large counties as well.

Realization that overdose deaths have been growing in a predictable exponential pattern for 37 years invites speculation that this pattern will continue into the near future. Based on 37 years of epidemic growth with a tight fit (R^2^ = 0.99) to an exponential curve, by simple extrapolation of that curve we forecast that in the next five years approximately 300,000 new drug overdose deaths can be expected to occur in the USA, if the curve is not deflected. Because this same exponential growth pattern holds for states and large counties, similar forecasts can be made for these jurisdictional levels, too. Such forecasts may be useful in planning and allocations of resources

This straightforward curve fitting approach can be used to plot the expected trajectory of the epidemic in any given jurisdiction, which can serve as a baseline for evaluation of the impact of interventional programs. By comparing expected and observed trajectories across several jurisdictions, it should be possible to identify those jurisdictions that have had the greatest success in deflecting the expected epidemic curve.

We were surprised to find that counts of overdose deaths in the USA have been following a clear exponential growth pattern since at least 1979. Throughout this 37 year interval there have been major fluctuations in the drugs most commonly used, as well as changes in legal prescribing, illicit supply, substance use treatment, and law enforcement. Our observation that this continuous exponential growth pattern began well before the onset of the current opioid epidemic suggests that deeper sociological, economic, and technological factors may be operative.

